# Synonymous mutations can alter protein dimerization through localized interface misfolding involving self-entanglements

**DOI:** 10.1101/2021.10.26.465867

**Authors:** Lan Pham Dang, Daniel Allen Nissley, Ian Sitarik, Quyen Vu Van, Yang Jiang, Mai Suan Li, Edward P. O’Brien

## Abstract

Synonymous mutations in messenger RNAs (mRNAs) can reduce protein-protein binding affinities by more than half despite leaving the protein’s amino acid sequence unaltered. Here, we use coarse-grain simulations of protein synthesis, ejection from the ribosome, post-translational dynamics, and dimerization to understand how synonymous mutations can influence the dimerization of the two *E. coli* homodimers oligoribonuclease and ribonuclease T. We synthesize each protein from its wildtype, fastest- and slowest-translating synonymous mRNAs and calculate the ensemble-average interaction energy between the resulting dimers. We find, similar to experiments with other dimers, that oligoribonuclease’s dimerization is altered by synonymous mutations. Relative to wildtype, the dimer interaction energy becomes 4% and 10% stronger, respectively, when translated from its fastest- and slowest-translating mRNAs. Ribonuclease T dimerization, however, is insensitive to synonymous mutations. The structural and kinetic origin of these changes are misfolded states containing non-covalent lasso-entanglements, many of which structurally perturb the dimer interface, whose probability of occurrence depends on translation speed. Translation of the fast- and slow-translating mRNAs of oligoribonuclease decreases the population of these misfolded states relative to wildtype. For ribonuclease T, however, these misfolded populations are insensitive to synonymous mutations. Entanglements cause altered dimerization energies for oligoribonuclease as there is a significant association (odds ratio: 50) between non-native self-entanglements and weak-binding dimer conformations. These conclusions are independent of model resolution, as entangled structures persist in long-time-scale all-atom simulations. Thus, non-native changes in entanglement is a mechanism through which oligomer structure and function can be altered.

**SIGNIFICANCE STATEMENT:** Synonymous mutations affect a range of post-translational protein functions, including dimerization, without altering the amino acid sequence of the encoded protein. This suggests that proteins somehow retain a “memory” of their translation-elongation kinetics long after synthesis is complete. Here, we demonstrate that synonymous mutations can change the likelihood that nascent proteins misfold into self-entangled conformations. These self-entangled structures are similar to the native state but with key conformational perturbations that disrupt the dimer interface, reducing their ability to dimerize. Rearrangement of such self-entangled states to the native state is a slow process, offering a structural explanation for how translation-elongation kinetics can influence long-time-scale protein-protein binding affinities.

## INTRODUCTION

Oligomerization, the process of assembling multiple macromolecules into dimers and higher-order oligomers, is necessary for a majority of proteins to function^1^. These functional oligomeric assemblies require the correct type, number, conformational state, and orientation of each constituent protein monomer^1^. For example, the monomers composing the active tetrameric forms of β-galactosidase^2^ and hemoglobin^3^ do not function efficiently on their own. An analysis of 452 human enzymes found roughly one-third (141) to be monomeric, one-third to be homodimers (125), and the remaining third to be heterodimers or higher order oligomers^4^. Just as the native structures of proteins represent their minimum free energy structure at equilibrium, thermodynamics is also thought to dictate the structural ensemble of oligomeric complexes. From this thermodynamic perspective, the initial conditions and history associated with a system have no long-term effect on its behavior, meaning that the influence of translation-elongation kinetics is irrelevant.

Contrary to this prediction, experiments have revealed that changes to the speed of protein translation can perturb post-translational oligomerization and protein function over biologically long timescales, indicating a role of kinetics and changes in co-translational processes. For example, when the sub-optimal codon usage in the *frq* gene encoding the FRQ circadian clock protein in *N. crassa* is “optimized” by replacing rare codons with common synonymous codons that tend to be translated quicker, it binds 60% less to the WC-1 protein even after controlling for soluble expression level changes. This decrease in dimerization effectively abolishes *N. crassa’s* circadian rhythm measured over the course of multiple days^5^. Thus, synonymous mutations can change the structure and function of oligomers and cause phenotypic changes.

Recent studies^6,7^ have suggested a mechanism by which synonymous mutations can alter monomeric protein enzyme structure and function, and how these changes can persist in the presence of the proteostasis machinery – such as chaperones and the proteasome – that evolved to fix or remove misfolded proteins. These studies indicate that long-lived misfolded states are self-entangled, leading to reduced structure and function. Many of these entangled structures largely resemble the native state and thus can evade chaperones, avoid aggregation, and fail to be degraded, allowing them to remain soluble but less functional on timescales ranging from seconds to months or longer. The partitioning of nascent proteins into such soluble but self-entangled conformations has the potential to explain how changes to translation kinetics are able to disrupt oligomer formation for long time periods.

Here, we use coarse-grain and all-atom molecular dynamics simulations to understand the structural origin of altered dimerization when synonymous mutations are introduced into a protein’s mRNA template. Because FRQ is an intrinsically disordered protein^8^ whose binding interface and structure are unknown, we instead study the dimerization of two paralogous, globular cytosolic *E. coli* homodimers - oligoribonuclease and ribonuclease T - after synthesis from their wildtype, fastest-translating synonymous variant, and slowest-translating synonymous variant mRNA sequences. For ribonuclease T, which folds relatively quickly, the speed of translation has no discernible influence on its ability to dimerize. Oligoribonuclease’s dimerization, however, does depend on the mRNA variant from which it is synthesized. We find a molecular origin of this phenomenon, show the results are robust to changes in model resolution, and explain why the mechanism we identify is likely to be widespread across the proteome.

## METHODS

### Construction of coarse-grain protein and ribosome representations

We employ a previously published Gō-like coarse-grain methodology in which each amino acid is represented by a single interaction site^9–11^. Briefly, the potential energy of a configuration in this model is computed by the equation

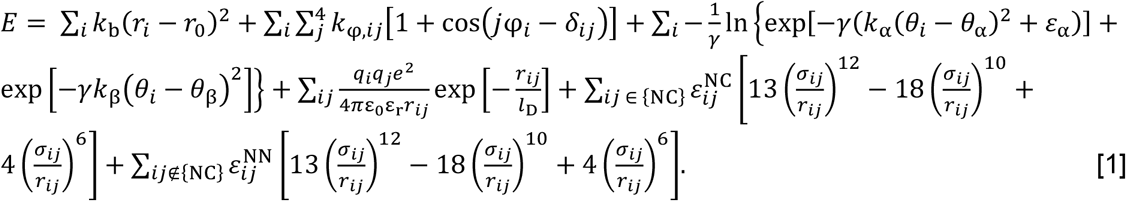

The summation terms in Eq. 1 represent, respectively, the contributions from C_α_-C_α_ virtual bonds, dihedral angles, bond angles, electrostatic interactions, Lennard-Jones-like native interactions, and repulsive non-native interactions to the total potential energy of the coarse-grain model. Bonds are treated using a harmonic potential and dihedral terms are computed as previously described^12^. The bond angle energy is computed using a double-well potential that can adopt angles representative of either α or β structures^13^. Electrostatics are treated using Debye-Hückel theory with a Debye screening length, *l*_D_, of 10 Å and a dielectric of 78.5. Lysine and arginine residues are assigned a charge of *q* = +1*e*, glutamate and aspartate *q* = −1*e*, and all other residue types *q* = 0^[10]^. Native and non-native interactions are computed using the 12-10-6 potential of Karanicolas and Brooks^12^. The minimum potential energy of a native contact is calculated as 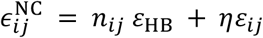. where *ε*_HB_ = 0.75 kcal/mol and *ε_i j_* represent the energy contributions arising from hydrogen bonding and van der Waals interactions between residues *i* and *j*, respectively, and *n_i j_* is the number of hydrogen bonds between residues *i* and *j. η* is a scaling factor that multiplicatively increases the values of *ε_i j_*, which are initially set based on the Betancourt-Thirumalai pairwise potential^14^. The collision diameter for a native contact between residues *i* and *j*, denoted *σ_i j_*, is set equal to the distance between the C_α_ atoms of the corresponding residues in the native-state crystal structure divided by 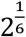. Values of *η* were determined based on a previously published training set to reproduce realistic protein domain stabilities (see below)^11^. Interactions between all pairs of residues not in the native contact list are computed with 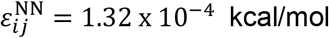 and collision diameters calculated using the Betancourt-Thirumalai algorithm^14^.

Synthesis simulations were conducted using a previously described protocol^9^ with a cutout of the ribosome exit tunnel and surface. Briefly, ribosomal RNA is represented with one bead for the ribose, phosphate, and pyrimidine bases and two beads for purine bases; ribosomal proteins are coarse-grained at C_α_-resolution as described for other proteins above^9,10^. The peptidyl-transferase center is placed at the origin of the CHARMM internal coordinate system, with the positive *x*-axis pointing down the ribosome exit tunnel towards its opening into the cytosol. All coarse-grain simulations were carried out with a Langevin thermostat set to 310 K, a 15-fs integration time step, and a friction coefficient of 0.050 ps^−1^.

### Parameterization of intra- and inter-monomer protein interactions

We set realistic intra- and inter-monomer energy scales in the coarse-grain models of PDB IDs 1YTA and 2IS3, which represent the homodimeric structures of the *E. coli* proteins oligoribonuclease and ribonuclease T, respectively, by tuning the scaling factor *η* separately for intra- and inter-monomer native contacts. Missing heavy atoms and residues in the ribonuclease T structure were reconstructed based on default CHARMM topology and parameters and then locally minimized as previously described^9^ before construction of its coarse-grain representation. A coarse-grain monomer is considered to be reasonably stable if its fraction of native contacts, *Q*, is greater than the average 〈*Q*_kin_〉 = 0.69 determine from the training set^11^ for at least 98% of the simulation frames in each of three 1-μs simulations initiated from the native state reference structure. The minimum value of *η* that results in a stable model based on these criteria is selected for each monomer and interface. A value of *η*_intra_ = 1.359 was selected by this procedure for all intra-monomer contacts in both oligoribonuclease and ribonuclease T; inter-monomer contacts were scaled by *η*_inter_ = 1.507 and 1.235 for oligoribonuclease and ribonuclease T, respectively.

### Construction of mRNA protein templates for coarse-grain simulations

Wildtype mRNA sequences for oligoribonuclease and ribonuclease T were obtained from NCBI assembly eschColi_K12 using the University of California Santa Cruz microbe table browser (http://microbes.ucsc.edu/). Codon translation rates are taken from the Fluitt–Viljoen^15^ model for *E. coli*, rescaled to produce an overall average elongation rate of 20 aa/s, and then further adjusted to account for the accelerated timescale of dynamic processes in our coarse-grain model. When rescaled in this way, the translation times from the Fluitt-Viljoen model have a mean of 12.6 ns or 840,000 integration time steps of 15 fs duration (see Supplementary Table 7 of Ref. 9). Predicted fastest- and slowest-translating synonymous mutant mRNAs were generated for oligoribonuclease and ribonuclease T by replaced each codon in their wildtype sequences with the codon predicted by the Fluitt-Viljoen model to be fastest- or slowest-translating, respectively. The average *in silico* translation times for the codons within the wildtype, fast-translating mutant, and slow-translating mutant sequences of oligoribonuclease are 10.6, 7.0, and 20.8 ns, respectively. Average translation times for the ribonuclease T wildtype, fast-translating mutant, and slow-translating mutant mRNAs are 12.7, 7.2, and 22.4 ns, respectively.

### Coarse-grain simulations of monomer synthesis, ejection, and post-translational dynamics

One hundred statistically independent continuous synthesis simulations were performed as previously described for each monomer of each protein and for each mRNA (*i.e*., 200 trajectories per protein, 100 for Monomer A and 100 for Monomer B, for each of the wildtype, fast-translating mutant, and slow-translating mutant mRNAs)^9^. In these simulations, a coarse-grain cutout of the ribosome exit tunnel and 50S surface consisting of 3,800 interaction sites is explicitly represented (Figure 1a, b, and c panel 1). The dwell time at a particular nascent chain length *k* was randomly selected from an exponential distribution with a mean equal to the average decoding time of the *k* + 1 codon (*i.e*., the time to decode the codon in the ribosomal A-site). After synthesis was completed, the harmonic restraint on the C-terminal bead representing the covalent bond between the nascent protein and the P-site tRNA is removed, allowing ejection of the nascent protein from the exit tunnel (Figure 1c, panel 2). Simulations of nascent protein ejection were run until the C-terminal residue reached an *x*-coordinate of 100 Å or greater in the internal CHARMM coordinate system. After ejection, the ribosome representation was deleted and 5 μs of post-translational dynamics simulated for each trajectory (Figure 1c, panel 3).

**Figure 1.**
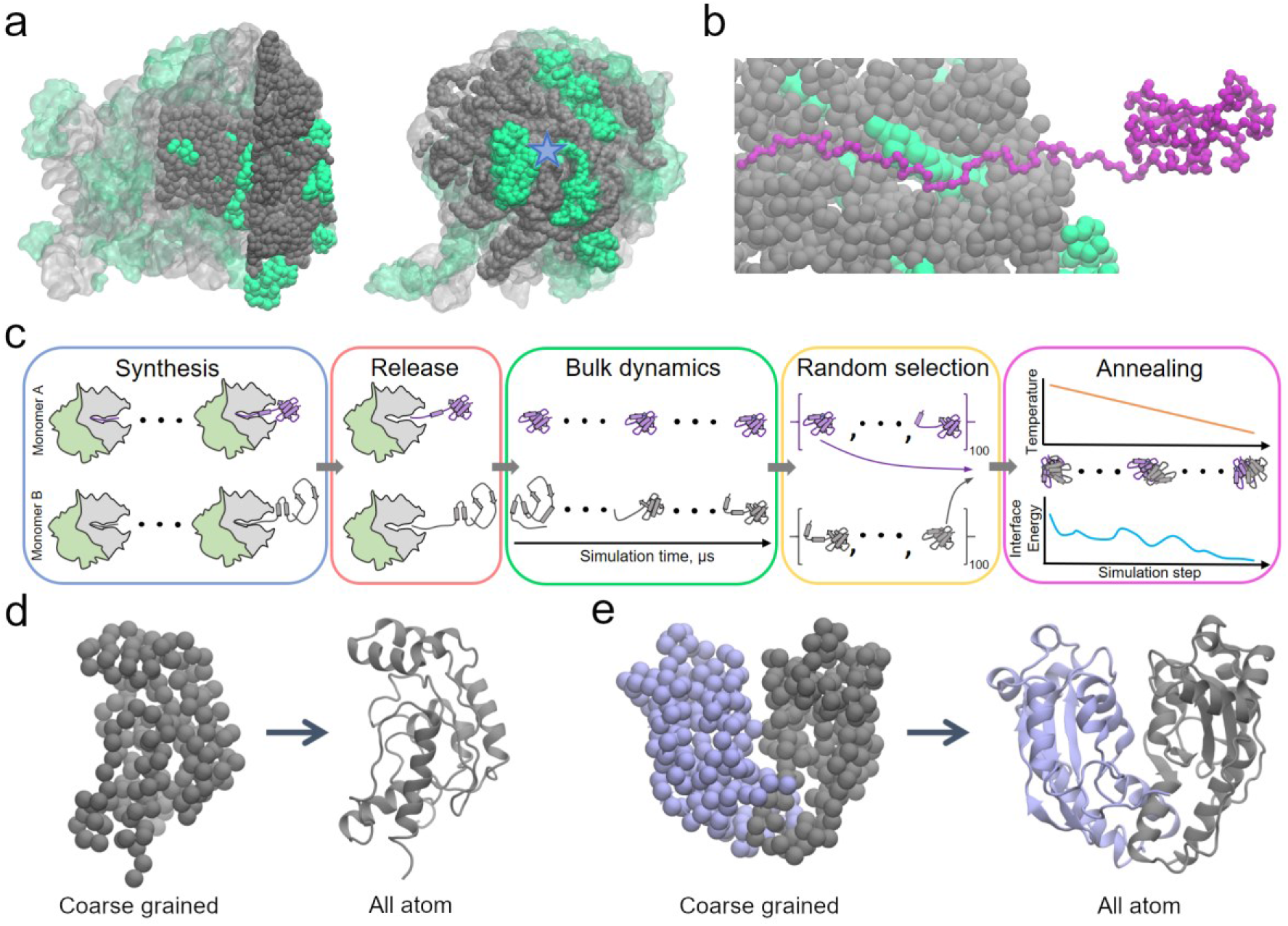
Simulating protein dimerization and entanglement at multiple resolutions. (a) Side (left) and top (right) views of the coarse-grained 50S *E. coli* ribosome cutout (filled spheres) used in our simulations superimposed over the entire all-atom 50S subunit (transparent) from PDB ID 3R8T. Ribosomal RNA and protein are displayed in grey and green, respectively. The approximate location of the ribosome exit tunnel is indicated by a blue star in the top view. (b) Side view of a 181-residue oligoribonuclease ribosome nascent chain complex just prior to its release from the ribosome. Note that the side of the exit tunnel was made cut away for visualization only. (c) Schematic of simulation protocol. One hundred nascent protein conformations are generated for Monomer A (purple) and Monomer B (grey). Each monomer is synthesized one amino acid at a time using coarse-grain protein and ribosome models (represented here by the purple/grey lines and green/grey shapes, respectively). After synthesis, the monomer is released from the ribosome and its bulk dynamics then simulated for 5 μs. Random combinations of the final structures from bulk dynamics are then selected from the sets of 100 Monomer A and 100 Monomer B trajectories and their lowest-energy dimer configurations determined by temperature annealing. (d) Initial coarse grain and resulting all-atom structures of oligoribonuclease monomer before and after back-mapping. (e) Same as (d) but for a dimeric oligoribonuclease structure.

### Computing the average interface interaction energy between monomers

Two hundred pairs of monomer structures were randomly selected from the 100 final conformations of Monomer A and 100 final conformations of Monomer B obtained after 5 μs of post-translational dynamics for a given protein and synonymous mRNA (Figure 1c, Panel 4). To generate dimer structures, the random monomer pairs were first aligned to the crystal structure coordinates based on interface residue locations only. Steric clashes were then resolved by an iterative procedure to identify the lowest-energy dimer structure. In this procedure, the interaction energy between the two monomers is first calculated in CHARMM. If the energy is positive (*i.e*., non-attractive), the Monomer B structure is translated 0.5 Å away from the Monomer A structure along the vector connecting their interface centers of mass. This procedure is terminated when the interaction energy is found to be less than or equal to zero. This conformation is then used as the initial condition for annealing simulations (Figure 1c, Panel 5). During annealing, the dimer structure is cooled from 310 to 0 K in 5-K increments. At each temperature, 150 ps of Langevin dynamics is simulated to allow for structural rearrangement of the interface. During annealing, a harmonic root mean square deviation restraint with force constant 5 kcal/[mol x Å^2^] is applied to each monomer to maintain them in their initial conformations. Five hundred independent annealing simulations were run for each pair of randomly selected monomers and the structure with the lowest interface interaction energy after annealing selected. The average interaction energy between monomers generated for a particular protein and mRNA reported in Figure 2 is computed as the mean of these 200 lowest-energy values found from the sets of 500 annealing simulations for each of the 100 random dimer structures.

**Figure 2.**
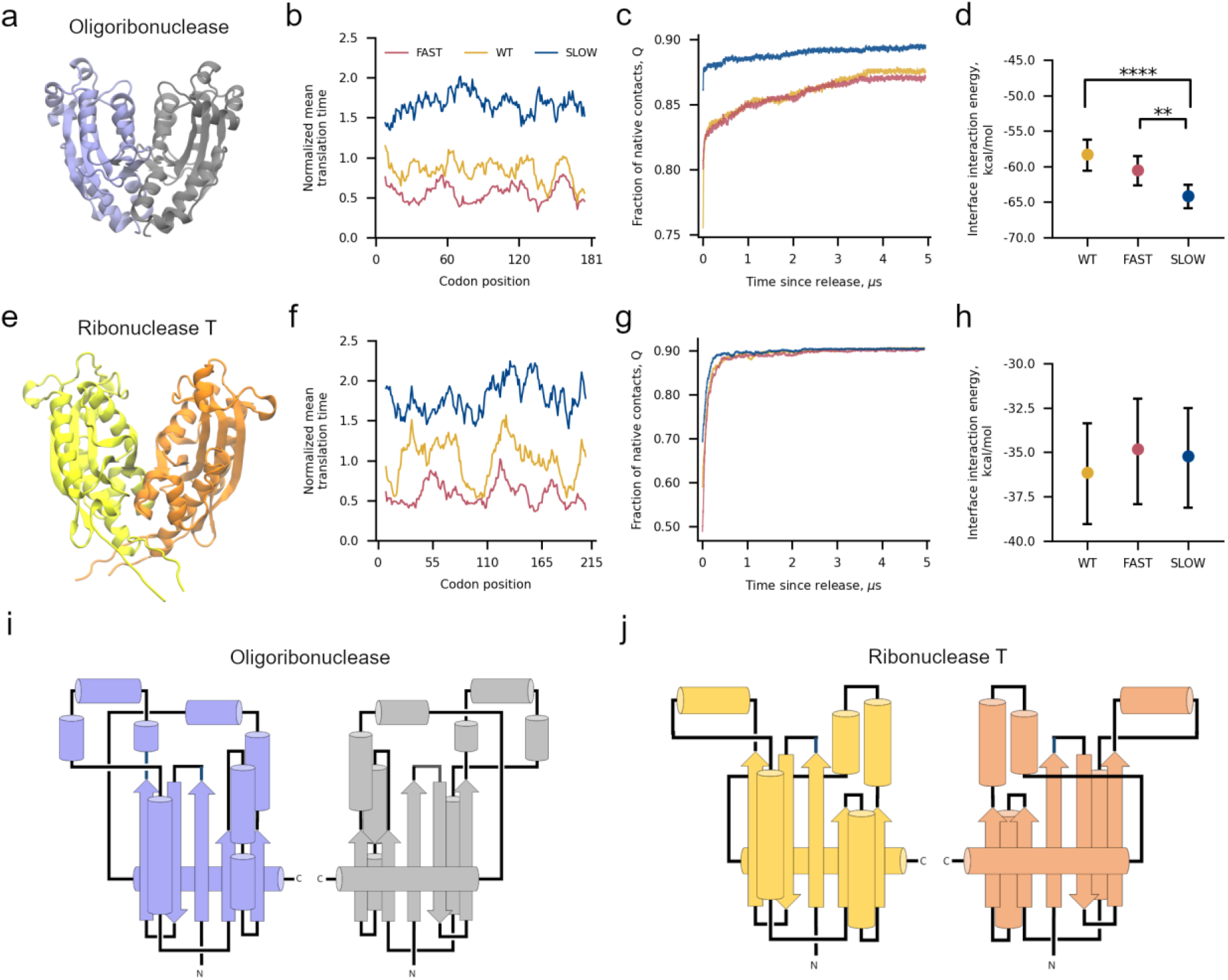
Altering translation kinetics affects the binding affinity of the oligoribonuclease homodimer. (a) 3D structure of oligoribonuclease from PDB ID 1YTA with Monomers A and B colored light purple and grey, respectively. (b) Normalized mean translation time of codon positions within the fast-translating mutant (FAST, red), wildtype (WT, yellow), and slow-translating mutant (SLOW, blue) mRNA sequences used for oligoribonuclease simulations smoothed with a 15-codon moving average. (c) Average fraction of native contacts as a function of time since oligoribonuclease’s release from the ribosome computed over all 200 trajectories (100 Monomer A + 100 Monomer B) for each mRNA. (d) Average interface interaction energy between Monomers A and B computed over 200 different random pairs of monomers after annealing as described in Methods and Figure 1c. Error bars are 95% confidence intervals computed from bootstrapping 10^6^ times. Brackets and asterisks indicate statistical significance of comparisons between means determined from permutation tests with 10^6^ samples. (e) 3D structure of ribonuclease T from PDB ID 2IS3 with monomers A and B colored yellow and orange, respectively. (f) Normalized mean translation time of codon positions in ribonuclease T mRNAs used in our simulations. (g) Fraction of native contacts versus time computed from 200 Ribonuclease T trajectories for each mRNA template used. (h) Same as (d) but for interactions between monomers of ribonuclease T. (i) Schematic representation of the native homodimeric structure of oligoribonuclease. (j) Same as (i) but for ribonuclease T.

### Extrapolation of folding times after release from the ribosome

A given trajectory of monomeric oligoribonuclease or ribonuclease T is considered to fold after its release from the ribosome when its *Q* first reaches ≥0.69 and remains ≥0.69 for at least 750 ps^[11]^. Based on this definition of folding, we computed the survival probability of the unfolded state of each protein as a function of time, denoted *S*_U_(*t*). The resulting time series were then fit to the double-exponential equation *S*_U_(*t*) = *f*_1_exp(−*k*_1_*t*) + *f*_2_exp(−*k*_2_*t*) with *f*_1_ + *f*_2_ ≡ 1. This fit equation represents a kinetic scheme in which folding proceeds through an obligate intermediate before proceeding irreversibly to the native state. The folding times of the two kinetic phases are computed as *τ*_1_ = 1/*k*_1_ and *τ*_2_ = 1/*k*_2_, with the larger of these two times determining the overall timescale of the folding process. The results of this fitting procedure for oligoribonuclease and ribonuclease T are summarized in Figure S1 and the resulting fit parameters are listed in Table S1.

### Identifying changes in entanglement and the residues involved in entanglements

To detect non-covalent lasso-like entanglements we exploit the numerically invariant linking numbers^16^, which describe the linking between two closed loops in three-dimensional space. This procedure is a modified version of a protocol previously used to detect entanglement in coarse grain protein structures^6^. The first loop is composed of the peptide backbone connecting residues *i* and *j* that form a native contact in a given protein conformation. Formation of the native contact between *i* and *j* is considered to close this loop, even though there is no covalent bond between these two residues. Outside this loop is an N-terminal segment, composed of residues 5 through *i* − 4, and a C-terminal segment composed of residues *j* + 4 through *N* − 5 for oligoribonuclease. Similar terminal ranges were used for ribonuclease T, but with a 15-residue terminal offset instead of 5 to address transient virtual entanglements caused by the long flexible tails of ribonuclease T. These two segments represent open curves when virtual bonds are drawn between the termini and the base of the loop. We can characterize the entanglement of the tails with each backbone loop formed by native contacts with 2 partial linking numbers denoted *g*_N_ and *g*_C_. We use the partial Gaussian double integration method of Baiesi and co-workers^17^ to calculate these partial linking numbers. For a given structure of an *N*-residue protein, with a native contact present at residues (*i, j*), the coordinates ***R**_l_* and the gradient *d**R**_l_* of the point *l* on the curves were calculated as

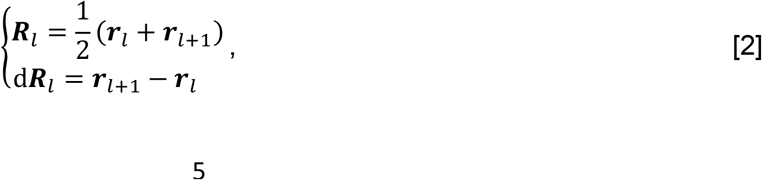

where ***r**_l_* is the coordinates of the C_α_ atom in residue *l*. The linking numbers *g*_N_(*i, j*) and *g*_C_(*i, j*) were calculated as

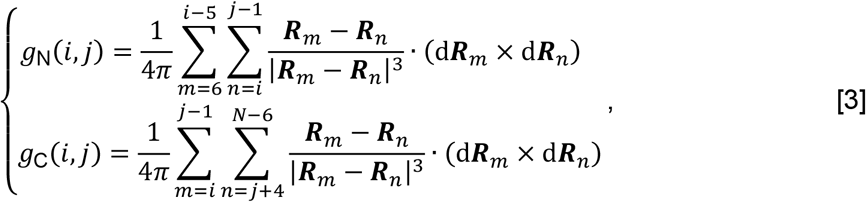

where we exclude the first 5 residues of the N-terminal curve, last 5 residues of the C-terminal curve, and 4 residues before and after the native contact to eliminate the error introduced by both the high flexibility and contiguity of the termini and trivial entanglements in local structure. The above integrations yield two non-integer values, and the total linking number for a native contact (*i, j*) was therefore estimated as

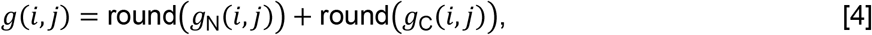

Comparing the absolute value of the total linking number for a native contact (*i, j*) to that of a reference state allows us to detect a gain or loss of linking between the backbone trace loop and the terminal open curves as well as any switches in chirality. Therefore, there are six change in linking cases we should consider when using this approach to quantify entanglement (see Table S1 of Ref. 6).

The N- and C-terminal threading locations, *g*_N|C_(*i, j, r*), of the most complex non-native entanglement is identified by first finding the native contact (*i, j*) where the total linking number is the equal to the global maximum of the set of all total linking numbers for the protein, *g*(*i, j*) = MAX[*g*(*i, j*)], and at which there was a change of entanglement detected. Second, using a sliding window of size 5, we find the terminal residues *r* where *g*_N|C_(*i, j, r*) = MAX[*g*_N|C_(*i, j, r*)] for the respective terminiThis method identifies residues involved in the change of entanglement when the window size is changed from 3 to 20 residues in width (Figure S2). A window size of 5 residues was chosen by examining ten random entangled structures and determining the minimal number of residues that can be visually distinguished as participating into the entanglement. Identifying the set of residues involved in the change of entanglement allows us to discern if the location disrupts the interface by examining the intersection of the set of interface residues with this set. An entanglement is considered to occur at the interface if any entangled residues are also identified to be interface residues, where interface residues are defined as those residues in Monomer A within 4.5 Å of Monomer B or *vice versa*.

### Clustering of dimeric entangled structures

Considering an ensemble of dimeric structures that contains at least one intra-monomer change in entanglement (note that no inter-monomer entanglements were observed) we separate them into clusters by examining the intersection between the sets of entangled residues in the two structures. Structures were merged in a leader algorithm^18^ style where the leader challenge is as follows:

1. Consider a leader superset of entangled residues *L* = {*A, B*} and a subordinate superset *s* = {*a, b*} where the sets *A, B, a, b* are the sets of residues in either monomer A(a) or B(b) that are involved in the entanglement.
2. Calculate the intersections *I_Aa_* = *A* ∩ *a*, *I_Bb_* = *B* ∩ *b*, and *I_Ba_* = *B* ∩ *a*
3. If |*I_Aa_*| & |*I_Bb_*| > 0 or |*I_Ab_*| & |*I_Ba_*| > 0, the subordinate passes the challenge and becomes part of the leader group. Otherwise, the challenge is failed and the search continues. If a subordinate fails the challenge of every current leader it become a new leader.

This approach ensures both identical and domain swapped entangled dimers are merged into the same state reflecting the homodimer nature of oligoribonuclease and ribonuclease T. The three most-populated (D1, D2, and D3) and the two lowest-energy (D4 and D5) dimeric entangled states of oligoribonuclease were selected for back-mapping to atomistic resolution as described below. Three all-atom monomeric systems were also generated by selecting three entangled monomers from within these five dimeric starting structures. (see Supplementary Data File 1).

### Back-mapping of coarse-grain monomer and dimer structures to all-atom resolution

Coarse-grain interaction sites representing the side-chain center-of-mass were rebuilt near their corresponding C_α_ beads in the selected dimer structures based on the native-state all-atom conformation^9^. Energy minimization was then performed using a two-bead C_α_-C_β_ coarse-grain force field^19^ generated from the original all-atom PDB with all C_α_ positions restrained. The backbone and sidechain atoms were then rebuilt with PD2^20^ and Pulchra^21^, respectively. The final all-atom structure was obtained after a further energy minimization *in vacuo* with all C_α_ positions restrained using OpenMM^22^. Representative starting coarse-grain and ending all-atom structures for the oligoribonuclease monomer and dimer are provided in Figure 1d and e, respectively.

### All-atom simulations of entangled structures

The back-mapped protein was placed in a rectangular box with a minimum distance of 1 nm between the edge of the protein and the periodic boundary wall in all dimensions. The system was solvated in TIP3P^[23]^ water and neutralized by Na^+^ and Cl^−^ counter-ions before adding 0.15-M sodium chloride to mimic the salt concentration inside the cell^24^. We next minimized and equilibrated the system. First, 1 ns of dynamics was carried out in the NVT ensemble, followed by 1 ns of dynamics in the NPT ensemble with harmonic position restraint potential (spring constant *k* = 1000 kJ/(mol x nm^2^)) applied to all heavy atoms of protein to relax the environment with the temperature and pressure held at 310 K and 1 atm, respectively. To allow the protein to reach equilibrium in the all-atom model and maintain the coarse-grain structure, we performed a second NPT simulation for 1 ns with harmonic restraints (*k* = 1000 kJ/(mol x nm^2^)) applied to all C_α_ atoms. Finally, we ran production simulations for 500 ns for five dimer and three monomer structures with three statistically independent trajectories with different initial velocities generated from the Maxwell distribution for each starting structure. Simulations for one randomly selected dimer and monomer structure were extended to 1 μs with no qualitative change in results. Simulations were performed with GROMACS 2018^[25]^ using the AMBER99SB-ildn forcefield^26^. The particle mesh Ewald method^27^ was used to calculate the long-range electrostatic interactions beyond 1 nm. Van der Waals interactions were calculated with a cut-off distance of 1 nm. The Nose-Hoover thermostat^28,29^ and Parrinello-Rahman barostat^30^ were employed to maintain the temperature and pressure at 310 K and 1 atm, respectively. The LINCS^31^ algorithm was used to constrain all bonds involved hydrogen atoms. An integration time step of 2 fs was used for all simulations.

### Calculating odds ratios and associated significance metrics

The 200 post-annealing dimer structures generated for each protein and mRNA were labelled as strong or weak binding based on whether they had an interaction energy less than or equal to a threshold value selected from the cumulative distribution function of the interaction energy of the ensemble of annealed wildtype dimer structures. This threshold was initially set to the value at which the cumulative distribution function of interaction energy equals 5% and then increased to 95% in 10% increments. The number of structures containing non-native changes in entanglement was counted for each of the thresholds {5%, 15%, 25%, 35%, 45%, 55%, 65%, 75%, 85%, 95%}. A contingency table at each threshold was then generated where the two events are: (1) strong or weak binding and (2) the presence or absence of non-native entanglements. Odds ratios and *p*-values for the contingency tables constructed in this way (Tables S2 & S3) were computed in Python3 using the fisher_exact function in the SciPy stats module.

## RESULTS

### Synonymous mutations alter oligoribonuclease’s post-translational structural ensemble and ability to dimerize

To test if oligoribonuclease’s ability to dimerize is perturbed by synonymous mutations we simulated its synthesis from mRNAs corresponding to its wildtype coding sequence, a slow-translating synonymous mRNA sequence composed of non-optimal codons, and a fast-translating synonymous mRNA sequence composed of optimal codons (Figure 2a, b). After synthesis, we simulated the release of oligoribonuclease from the ribosome followed by 5 μs of post-translational dynamics (the equivalent of approximately 20 s in real time^11^). We find that oligoribonuclease exhibits structural differences in its post-translational folding dynamics dependent on whether it was translated from the wildtype, fast-translating, or slow-translating mRNA sequences (Figure 2c). The slow-translating mRNA produces a protein structural ensemble with a higher average fraction of native contacts (*Q* = 0.89, 95% CI: [0.87, 0.90], computed from bootstrapping 10^6^ times) relative to the wild-type mRNA (*Q* = 0.86, 95% CI: [0.85, 0.88], computed from bootstrapping) 5 μs after ribosome release. The ensemble of minimum energy dimeric structures from temperature annealing simulations (Figure 1c), reveals that this increase in native structure results in a more favorable dimer interface interaction energy of −64.1 kcal/mol (95% CI [−65.7, −62.4], computed from bootstrapping) for the slow mRNA products compared to dimers produced from the wildtype mRNA of −58.3 kcal/mol (95% CI [−60.3, −55.9], computed from bootstrapping). The difference between these two average interaction energies is significant (*p* = 2 x 10^−5^, computed from permutation test with 10^6^ samples). The average dimer interaction energy of −60.5 kcal/mol (95% CI: [−62.5, −58.4], computed from bootstrapping) is also different for proteins produced from the fast-translating mRNA (Figure 2d). These results demonstrate that oligoribonuclease’s post-translational dimerization affinity is modulated by changes in translation-elongation speed.

### Dimerization of ribonuclease T is not influenced by synonymous mutations

In contrast to oligoribonuclease, ribonuclease T’s post-translational structural ensemble (Figure 2 e-g) and dimerization interaction energy are not dependent on the mRNA that encodes it (Figure 2h). No statistically significant differences are found between the average interface interaction energies of the dimer ensembles generated from the wildtype, fast-translating, or slow-translating mRNAs.

### Oligoribonuclease but not ribonuclease T frequently populates self-entangled states that involve interface residues

To understand what translation-speed-dependent structural changes in oligoribonuclease cause altered dimer interaction energies we use the maximum intrachain contact entanglement^32^ metric, *g*(*i, j*) (Eq. 4). This metric identifies even subtly misfolded protein conformations. The value of *g*(*i, j*) represents the Gauss linking number between a closed loop formed by a backbone segment between residues *i* and *j* that form a native contact and the pseudo-closed loops formed by the N- and C-terminal segments of a protein^17^. This metric of protein structure is topologically invariant^33^ and describes how portions of the protein are linked together in space, allowing for the identification of misfolded states as a change in the Gauss linking number of specific native contacts relative to the native structure. Schematic illustrations of the different entanglement types can be viewed in Figure S3^7^.

We identified various entanglements within both the monomeric and dimeric ensembles of oligoribonuclease that cause disruptions relative to the native state (Figure 2i). Representative structures of two frequently occurring entanglements, in which either residues 96-102 or residues 125-129 are entangled, are displayed in Figure 3a-d, and schematic representations are shown in Figure 3e and f. Entanglement can occur in an isolated monomer (Figure 3a), in one monomer that forms part of a dimeric complex (Figure 3b), or in both monomers in a dimer (Figure 3c,d). Each of these entangled structures is highly native; for example, the entangled dimer structure shown in Figure 3c and d has ≤3-Å RMSD from the native state (Figure 3g). The high similarity of these entangled conformations to the native state suggests that they may remain soluble and evade proteostasis quality controls^6^. Despite their similarity to the native state, entanglements can structurally perturb the dimer interface. In the case of the entanglement of residues 125-129 in Monomer A, misplacement of a loop segment causes the disruption of a β-sheet that forms part of the dimer interface (Figure 3h). This suggests that changes in dimer interaction energy are structurally caused by entanglements perturbing the dimer binding interface.

**Figure 3.**
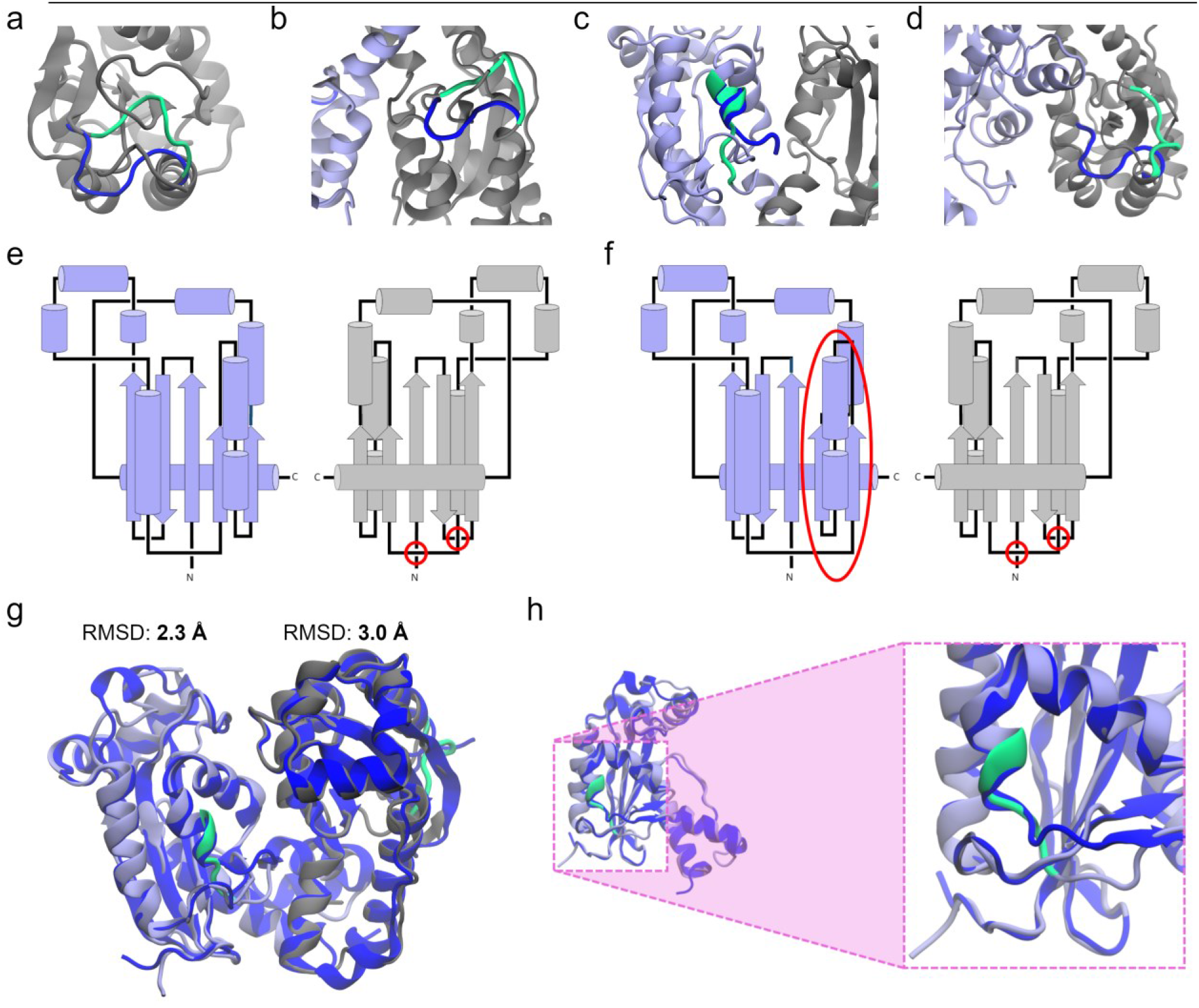
Entanglements in oligoribonuclease perturb its dimer interface. (a) Structure of oligoribonuclease Monomer B in which residues 96-102 are entangled (structure M3, see Supplementary Data File 1). (b) Same as (a) except in the context of a dimeric complex after annealing (structure D4). (c) Structure of oligoribonuclease dimer in which residues 125-129 of Monomer A and (d) residues 96-102 of Monomer B are entangled (structure D5). (e) Schematic showing the location of the entanglements shown in (a) and (b) in the context of the dimer structure. (f) Same as (e) except for the entanglements shown in (c) and (d). (g) Alignment of the Monomer A and Monomer B structures shown in (c) and (d) to the native state structure indicates they are overall native-like with ≤3-Å Root Mean Square Deviation (RMSD) from the crystal structure. (h) Left: interface view of the entangled structure of Monomer A from (c) aligned to its native state reference structure. Right: zoomed-in view of entangled residues 125-129 of Monomer A inserting below rather than above a loop segment, disrupting the formation of a β-sheet that forms part of the dimer interface. The correct β-sheet conformation is displayed in dark blue. Coarse-grain structures were back-mapped to all-atom resolution to generate visualizations.

### Synonymous mutations alter the population of self-entangled oligoribonuclease structures

To quantify the influence of synonymous mutations on the probability of entanglement within oligoribuclease and ribonuclease T we computed the fraction of monomers (out of 200 independent trajectories) and dimers (out of 100 annealed structures) that exhibit a non-native change in entanglement for both oligoribonuclease and ribonuclease T from their wild-type, fast- and slow-translating mRNAs (Figure 4). We find that ribonuclease T exhibits very little entanglement (<0.30) in both its monomeric and dimeric forms regardless of the translation-rate schedule used during its synthesis. Statistically significant differences are present depending on the translation schedule used (see, for example, Figure 4c); however, the magnitude of these population differences is small (less than 10%). This explains why ribonuclease T’s dimer interaction energy is insensitive to synonymous mutations – any corresponding population changes in misfolded states are quite modest and have little effect on this protein’s ability to dimerize.

**Figure 4.**
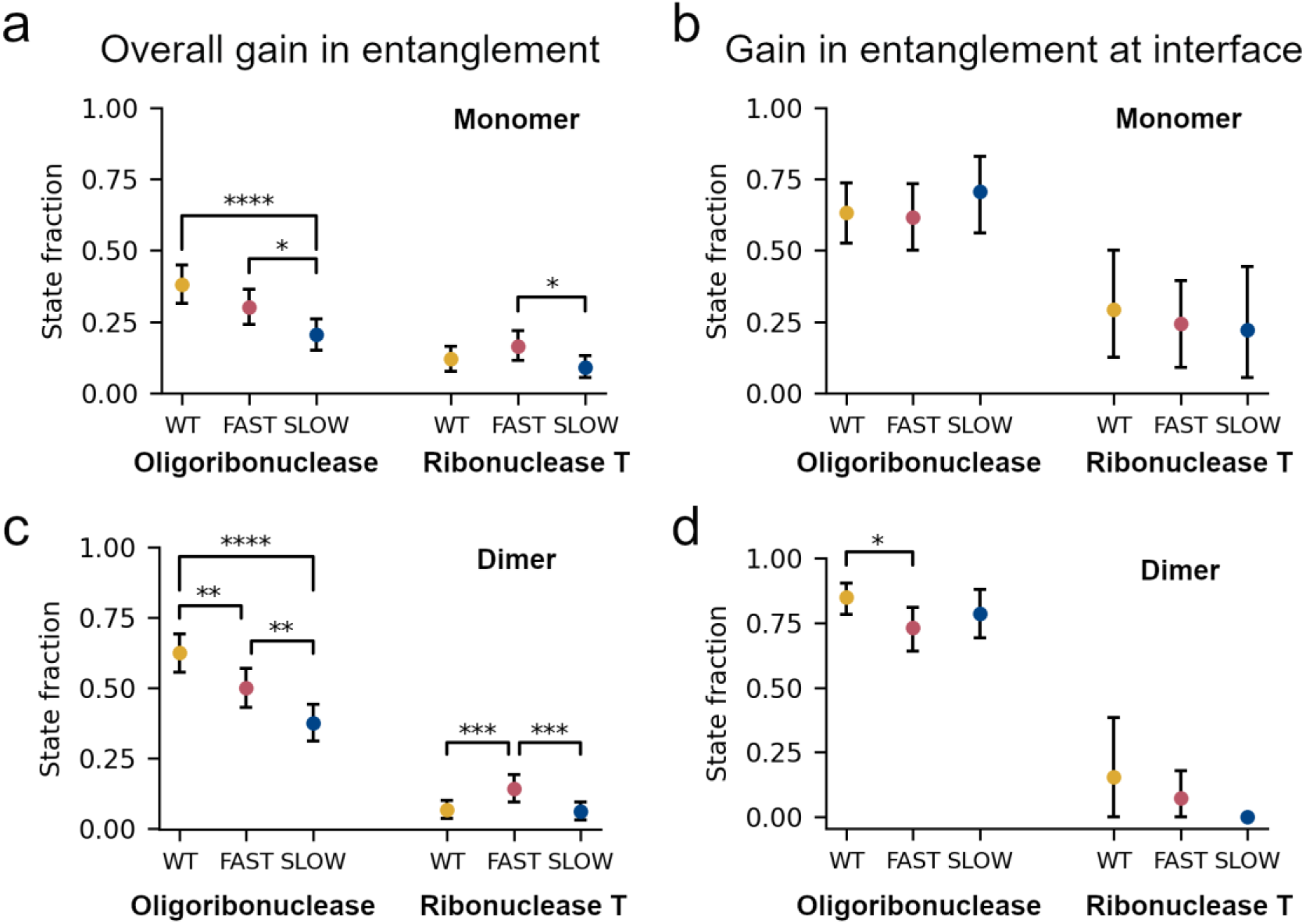
Changes in the population of self-entangled structures correlate with differences in dimer interaction energies. (a) Fraction of monomer structures of oligoribonuclease and ribonuclease T generated by coarse-grain synthesis and post-translational dynamics simulations using the wildtype (WT), fast-translating mutant (FAST), and slow-translating (SLOW) mutant mRNAs that have a gain in entanglement somewhere in their structure. (b) Same as (a) but limited to the specific set of entanglements involving residues at the dimer interface. (c) Same as (a) but computed for the dimer structures generated by annealing random pairs of monomers. (d) Same as (c) but limited to the specific set of entanglements involving interface residues. All error bars are 95% confidence intervals computed from bootstrapping 10^6^ times. Brackets and asterisks indicate the statistical significance of comparisons between means determined from permutation tests with 10^6^ samples.

In contrast, for oligoribonuclease, the population of conformations that display a non-native change in entanglement is larger in magnitude and more sensitive to changes in translation speed. For wildtype, fast, and slow synthesis the misfolded populations of states with an overall entanglement within the oligoribonuclease dimer are, respectively 0.63 (95% CI: [0.55, 0.69]), 0.50 (95% CI: [0.43, 0.57]), and 0.37 (95% CI: [0.31, 0.44], all computed by bootstrapping) (Figure 4c). This trend is consistent with the trend observed in the average dimer interaction energies (Figure 2d) where values of −58.3, −60.5, and −64.1 kcal/mol, respectively, are found for the wildtype, fast, and slow mRNA templates. These results suggest the hypothesis that changes in the population of entangled states with perturbed interfaces, arising from changes in translation speed, cause the binding affinity between monomers to be altered.

### Misfolded entangled states often involve the dimerization interface

Next, we quantitatively assessed the occurrence of entanglements at the dimer interface by computing the fractions of monomer and dimer states in which interface residues are entangled (see Methods and Figure 4). Ribonuclease T has low levels of entanglement at the interface, with state fraction less than 0.30 (Figure 4b and d), and there are no significant changes in entanglement when synonymous mutations are introduced. In contrast, oligoribonuclease displays much more frequent interface entanglement in both monomer and dimer structures (state fraction >0.50 in all cases, Figure 4b and d). Specifically, dimer structures with entanglements at the interface have state fractions of 0.85 (95% CI: [0.78, 0.90]), 0.73 (95% CI: [0.64, 0.81]), and 0.79 (95% CI: [0.69, 0.88], computed from bootstrapping) (Figure 4d). Thus, at least for oligoribonuclease, non-native entanglements commonly occur at the dimer interface.

### Entanglements greatly reduce the likelihood of strong dimer binding

To test whether entanglement is associated with decreased average binding energy between monomers, we create a two-by-two contingency table categorizing annealed dimer structures as strongly or weakly bound together and as having an entanglement present or not. A contingency table allows us to compute the conditional probability of these two events co-occurring, the odds ratio (effect size) that entanglement and binding are associated, and Fisher’s Exact test tells us whether the association is significant. Statistically significant odds ratios other than 1 would establish an association between these two phenomena.

To classify structures based on their binding energy, we define dimer complexes with interface interaction energy less than or equal to the 5^th^ percentile value from the wildtype interaction energy distribution to be strong binding and all others to be weak binding. Then, as a test of robustness, we systematically vary this threshold in increments of +10% up to 95% and compute the odds ratio and *p*-value for each splitting of the data using Fisher’s Exact test (Figure 5, contingency tables reported in Tables S2 and S3).

**Figure 5.**
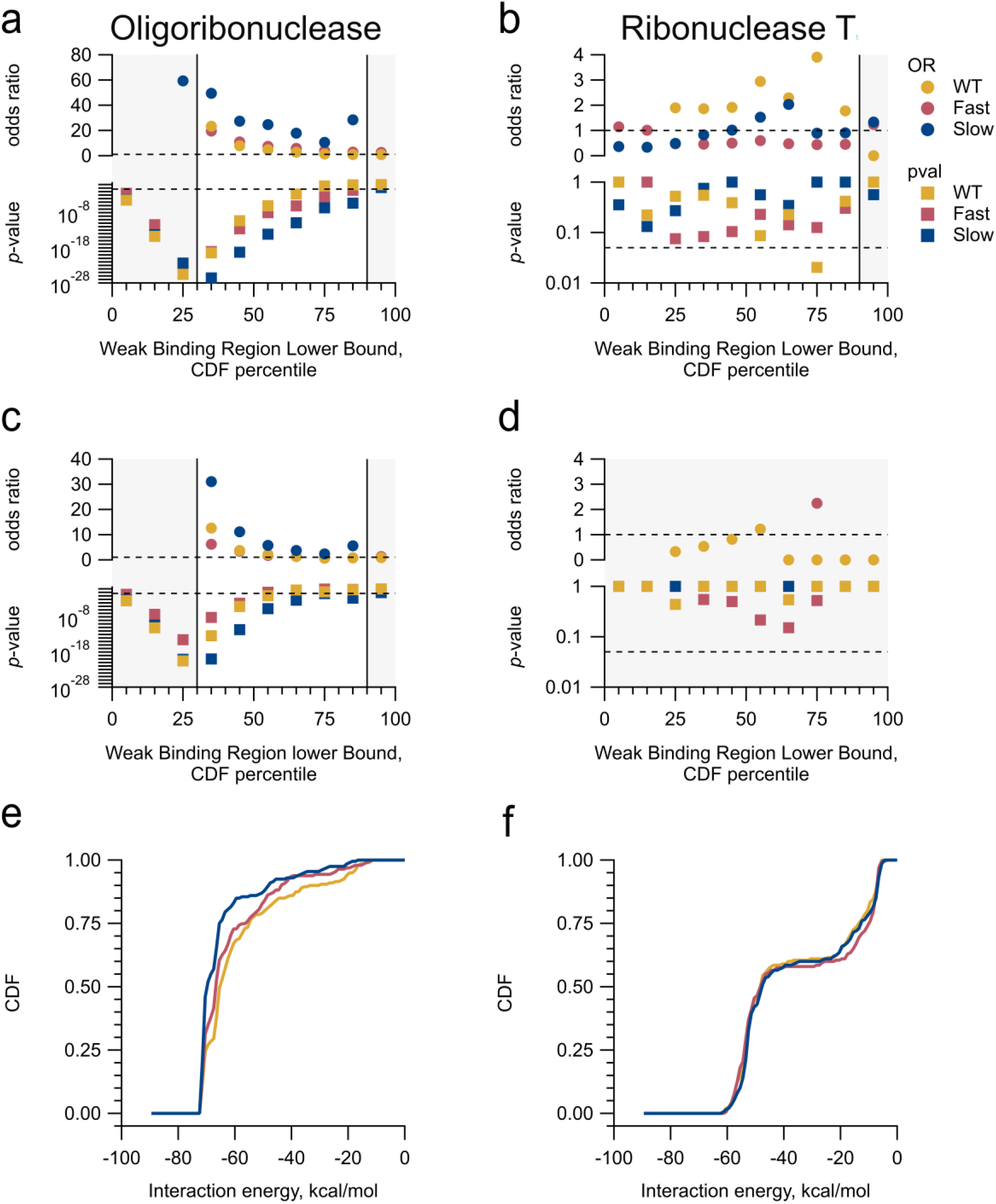
The presence of entanglements is strongly associated with weak dimer binding interaction energies. (a & b) Odds ratios and *p*-values resulting from Fisher’s Exact Test applied to contingency tables for oligoribonuclease and ribonuclease T. The events were defined as (1) the presence of any change in entanglement of the annealed dimer relative to a reference native state and (2) the interaction energy falls within the weak binding region. To test the robustness of the results, the upper bound of the strong binding region was swept from the 5^th^ percentile to the 95^th^ percentile of the appropriate WT distribution of interaction energies. Regions where the odds ratio is not well defined (*i.e*., 0 entries in the contingency table) are shaded in grey, and the dotted lines represent 1 and 0.05 on the odds ratio and *p*-value axes, respectively. (c & d) are the same as (a & b) but the first event is defined as the presence of entanglement at the interface (e & f) Cumulative distribution functions (CDFs) of the WT, FAST, and SLOW ensemble interaction energies.

For oligoribonuclease we find the odds ratio, where it is defined (unshaded portions of Figure 5a and b), is significantly greater than 1 for a range of thresholds. For example, at a 55% threshold, the odds ratio is 24.7 and is statistically significant (*p*-value = 7.6 x 10^−15^, Fischer’s Exact test; see Table S2) for the slow-translating mRNA variant. The magnitude of these odds ratios demonstrates there is a strong association between the presence of entanglements and the occurrence of dimers with weak dimerization energies.

In contrast, ribonuclease T has odds ratio’s that are not statistically different than one another at all threshold values (Figure 5b &d). This indicates, as inferred earlier, that the modest population of entangled structures for ribonuclease T has no association with strong or weak dimerization occurring.

### Entangled states are long-lived kinetic traps

Entangled states that are long lived can have long-term impacts on protein structure and function. Therefore, we quantified the lifetime of these states. While all monomers of ribonuclease T fold by 0.8 μs after release from the ribosome, some oligoribonuclease molecules fail to fold during the 5-μs post-translational simulations regardless of the mRNA template used (Figure S1). When synthesized from its wildtype, fast-translating, and slow-translating variants, respectively, 13% (95% CI [8%, 17%]), 14% (95% CI [9%, 19%]) and 10% (95% CI [6%, 14%], bootstrapping with 10^6^ iterations) of its monomers do not fold correctly (see Figure S1 and Methods). We used a kinetic curve-fitting procedure to estimate that these misfolded populations of oligoribonuclease require between 9 and 10 μs to fold (the equivalent of approximately 40 s of real time; Table S1). This indicates that these entangled states are kinetically trapped.

### Entanglements persist in all-atom molecular dynamics for up to one microsecond

To test whether our results generated using a C_α_ coarse-grain representation are resolution dependent we back-mapped representative entangled conformations of the oligoribonuclease dimer and monomer to atomistic resolution and simulated their aqueous dynamics for 500 ns. We ran three statistically independent trajectories for each of five different dimer starting structures and three different monomer structures selected to represent the entangled conformations most frequently populated and lowest in energy (see Methods and Supplementary Data File 1). In each case, the entanglement present in the coarse-grain model persists at all-atom resolution for the duration of the 500-ns simulation. In addition to these 500-ns simulations, we also extended the simulations for one randomly selected dimer and monomer structures to 1 μs and find the entangled states persist (data not shown). The time series of 〈*G*〉 and for four representative entanglements, one in a monomer and three in dimers, are displayed in Figure 6. Thus, the entangled structures we observe in our coarse-grained simulations can also be populated and persist in all-atom models.

**Figure 6.**
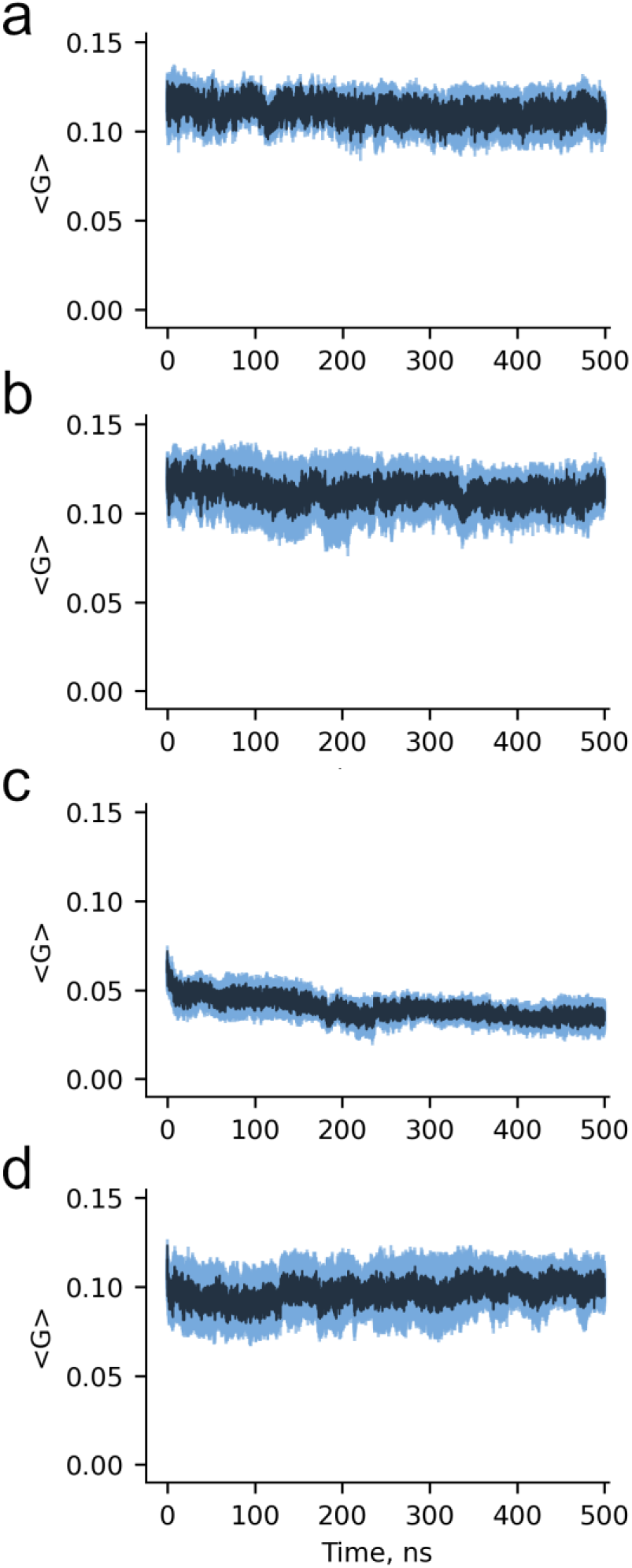
Entanglements in monomer and dimer structures of oligoribonuclease persist at all-atom resolution for 500 ns. (a) 〈*G*〉 (Eq. 4) as a function of simulation time computed over three all-atom trajectories each initiated from the same back-mapped entangled oligoribonuclease Monomer B structure (structure M3, see Supplementary Data File 1), in which residues 96-102 are entangled. The blue shaded region indicates the standard error of the average over the three trajectories. (b) Same as (a) except computed for dimer structure D4, in which residues 96-102 of Monomer B are entangled. The time series of 〈*G*〉 for Monomer A is not shown as it fluctuates around zero. (c) Same as (a) except for Monomer A of dimer structure D5, in which residues 125-129 are entangled. (d) Same as (a) except for Monomer B of dimer structure D5, in which residues 96-102 are entangled.

## DISCUSSION

Our results provide a structural explanation for how changes in translation speed induced by synonymous mutations can alter the ability of soluble proteins to dimerize over long time scales. For the homodimer oligoribonuclease, synonymous mutations change the population of protein molecules that partition into soluble, misfolded, self-entangled conformations. These entangled conformations – greater than 75% of which have an entanglement at the dimer interface – weaken the ensemble-averaged binding energy between the monomers over long time scales. In contrast, ribonuclease T is largely insensitive to synonymous mutations that alter translation speed, with far fewer states with entangled dimer interfaces (≤18%) populated and those that are entangled exhibiting little population dependence on translation speed. Thus, ribonuclease T’s dimer binding energy does not change with the introduction of synonymous mutations.

A commonly held assumption in the nascent protein folding field is that slower translation will result in more accurate protein folding. Therefore, one would predict that any changes in dimer interaction energy would follow the trend that the slow-translating mRNA will result in the strongest binding, followed by the wild-type and fast-variant mRNAs of oligoribonuclease. This is not what we observe – we find, respectively, that slow, fast, and wild-type mRNA variants result in increasingly weaker dimer affinities. This result is explained by both kinetic and simulation models showing the influence of translation kinetics on co-translational protein folding is, for some proteins, non-monotonic. Faster translation can result in an increased yield of correctly folded protein by translating quickly through protein segments that are prone to misfolding^34,35^.

Oligomer assembly can begin early in the life of a protein, with some nascent chains co-translationally dimerizing between adjacent ribosomes^1^. It is unknown how many different proteins engage in such co-translational assembly. Therefore, in this study we chose to consider the dimerization of nascent proteins after their release from the ribosome. Additionally, the motivating experiments on FRQ assessed only post- and not co-translational dimerization. Based on our results, we speculate that co-translational interface interaction energies are likely to follow similar mechanisms as we have identified in this post-translational study. Investigating how synonymous mutations influence co-translational dimerization is an interesting avenue for future research.

In summary, our results indicate that, at least for some proteins, synonymous mutations can modulate the amount of nascent protein that misfolds into soluble, non-native self-entangled conformations with reduced dimer interface interaction energies. For oligoribonuclease, slowing down or speeding up translation relative to its wildtype translation schedule leads to a reduction in entanglement, especially at the interface, leading to more stable dimers on average. In contrast, ribonuclease T folds quickly and is less prone to misfolding, and is therefore largely unaffected by synonymous mutations. Taken in combination with a recent large-scale study of entanglement in the *E. coli* proteome^6^ and an in-depth analysis of the influence of entanglement on enzymatic activity^7^, our results support the view that near-native, entangled misfolded states are likely to be a common phenomenon that influences a wide range of protein functions.

## DATA AVAILABILITY STATEMENT

Coarse-grain and all-atom molecular dynamics simulation data is available upon request. Codes used for back mapping to all-atom resolution and detection of self-entanglements are available at: https://github.com/orgs/obrien-lab/

## Supplementary Information

**Figure S1.**
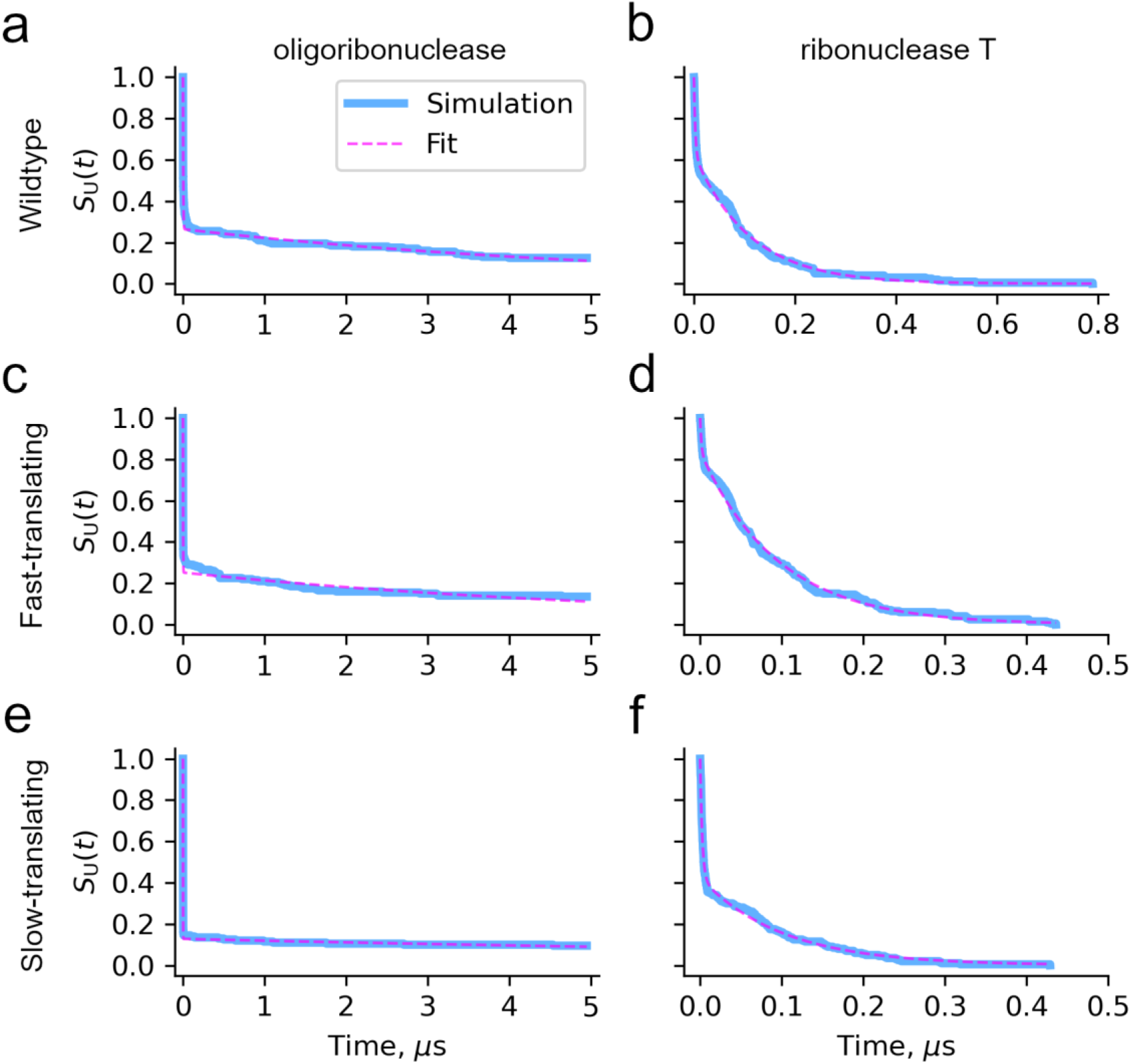
Calculation of post-translational folding timescales with curve fitting in Python. (a) Survival probability of the unfolded state as a function of time since release from the ribosome (blue) and fit to the double-exponential equation *S*_U_(*t*) = *f*_1_exp(−*k*_1_*t*) + *f*_2_exp(−*k*_2_*t*) (magenta dashed line, see Methods) for oligoribonuclease translated from its wildtype mRNA. (b) Same as (a) but for ribonuclease T wildtype mRNA simulations. (c) *S*_U_(*t*) and fit for oligoribonuclease fast-translating mRNA simulations. (d) Same as (c) but for ribonuclease T fast-translating mRNA simulations. (e) *S*_U_(*t*) and fit for oligoribonuclease slow-translating mRNA simulations. (f) Same as (e) but for ribonuclease T slow-translating mRNA simulations.

**Figure S2:**
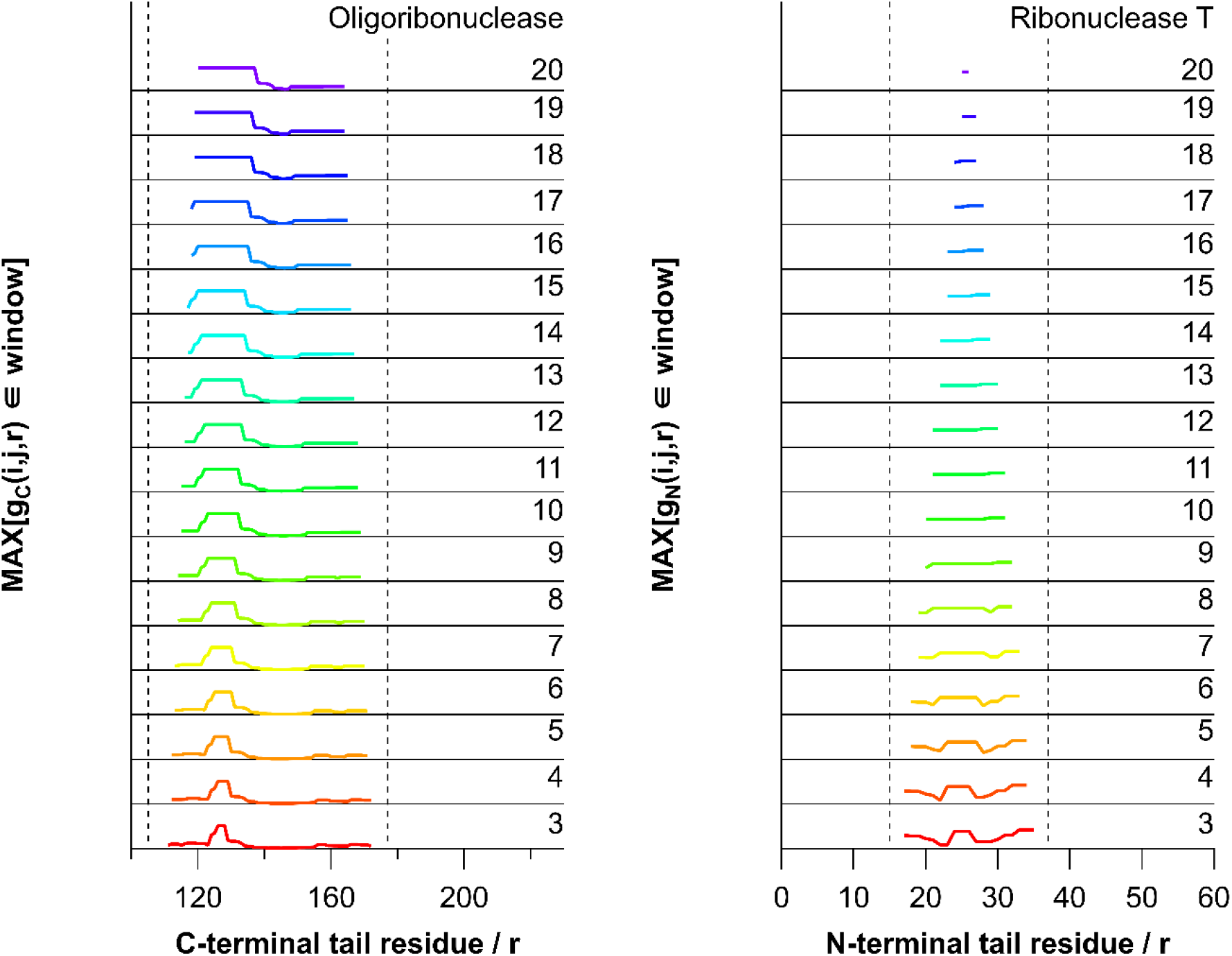
Example robustness analysis of the metric used to determine the residues involved in a change in entanglement. (left) raw trace of the maximal value of the C-terminal linking number for a structure in the oligoribonuclease wildtype ensemble. Window size is indicated by number of residues on the right axis. When the trace reaches a maximum the residues along the *x*-axis involved at the entanglement site. Dotted lines indicate the boundary of residues considered in the tail. (right) same as the (left) but for a structure in the ribonuclease T ensemble that has a predominate N-terminal tail threading of the loop instead. The residues involved in entanglement identified are consistent with those identified visually.

**Figure S3.**
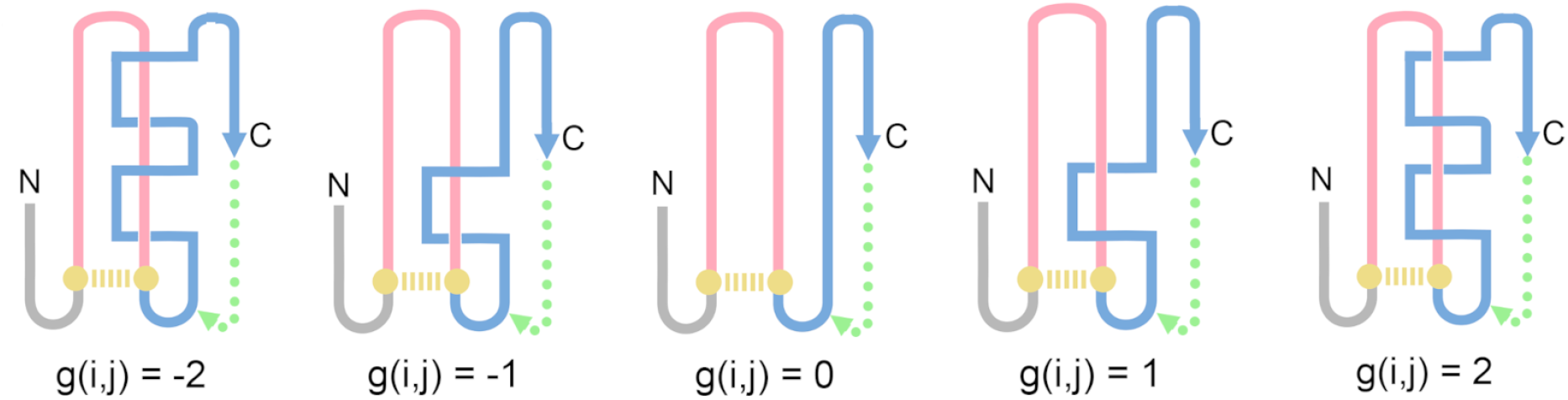
(a) Schematic of how misfolding can be detected by examining the change in the Gauss linking number *g*(*i, j*) (Eq. 4) for a closed loop (pink) formed by a given native contact between residues *i* and *j* (gold) and the C-termini (cyan) and the pseudo-closed loops formed by the flanking C termini (blue) and a virtual loop closure (dashed green line). The N-termini can also be used in the same manner.

**Table S1.** Kinetic fitting parameters to the equation *S*_U_(*t*) = *f*_1_exp(−*k*_1_*t*) + *f*_2_exp(−*k*_2_*t*) for oligoribonuclease and ribonuclease T post-translational folding time courses in Figure S1.

| Protein | mRNA | $f_1$ | $k_1, \mu\text{s}^{-1}$ | $\tau_1, \mu\text{s}$ | $f_2$ | $k_2, \mu\text{s}^{-1}$ | $\tau_2, \mu\text{s}$ | Pearson $R^2$ |
| --- | --- | --- | --- | --- | --- | --- | --- | --- |
| oligoribonuclease | wildtype | 0.26 | $1.74 \times 10^{-1}$ | 5.75 | 0.74 | $2.89 \times 10^2$ | $3.46 \times 10^{-3}$ | 0.95 |
| | fast | 0.25 | $1.65 \times 10^{-1}$ | 6.06 | 0.75 | $6.26 \times 10^2$ | $1.60 \times 10^{-3}$ | 0.86 |
| | slow | 0.13 | $7.10 \times 10^{-2}$ | $1.41 \times 10^1$ | 0.87 | $1.19 \times 10^3$ | $8.40 \times 10^{-4}$ | 0.86 |
| ribonuclease T | wildtype | 0.63 | 9.07 | $1.10 \times 10^{-1}$ | 0.37 | $3.85 \times 10^2$ | $2.60 \times 10^{-3}$ | 0.99 |
| | fast | 0.84 | $1.05 \times 10^1$ | $9.52 \times 10^{-2}$ | 0.16 | $6.56 \times 10^2$ | $1.52 \times 10^{-3}$ | 1.00 |
| | slow | 0.42 | 9.88 | $1.01 \times 10^{-1}$ | 0.58 | $3.91 \times 10^2$ | $2.56 \times 10^{-3}$ | 0.99 |

**Table S2.**
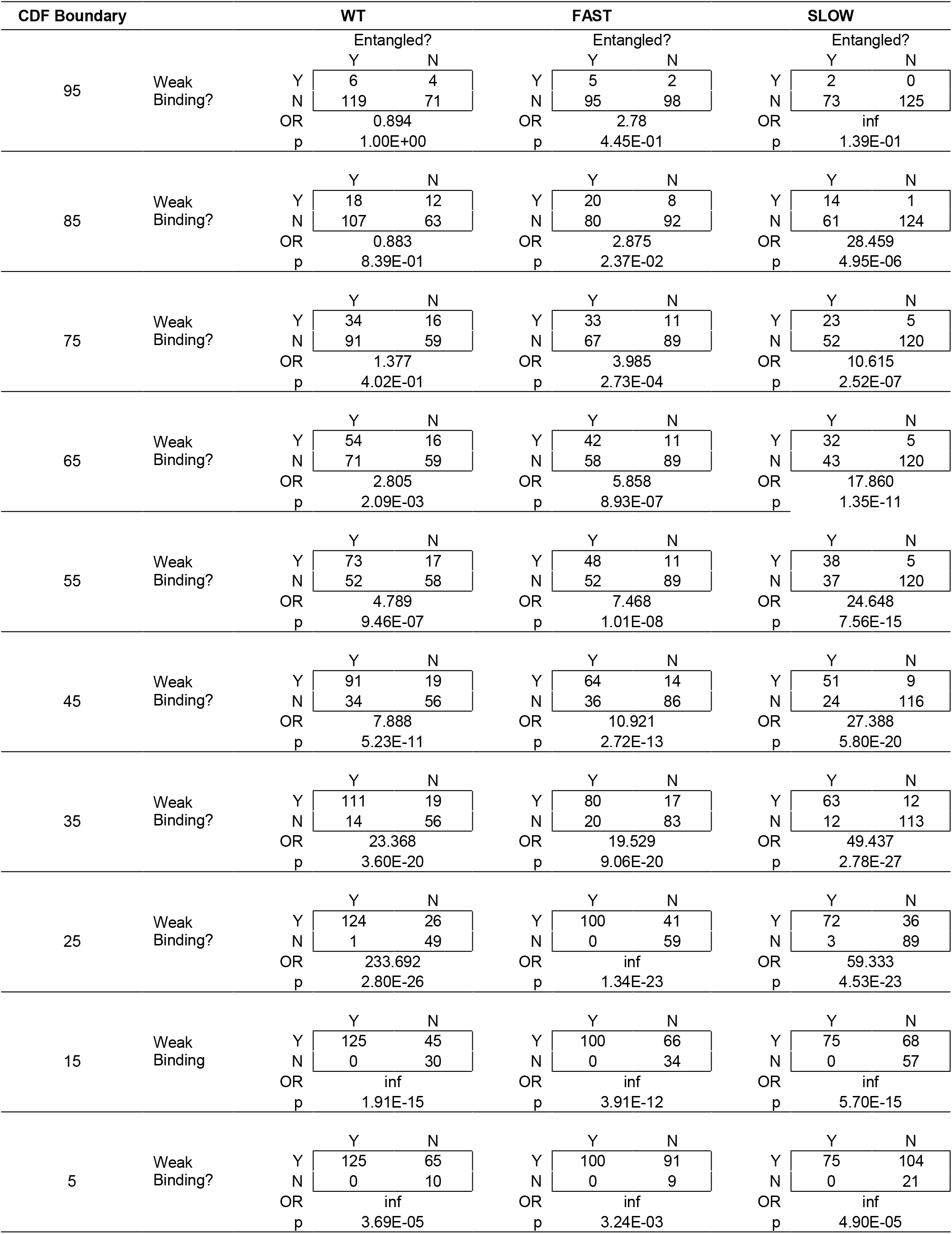
Oligoribonuclease entanglement & strong binding region contingency tables

**Table S3.**
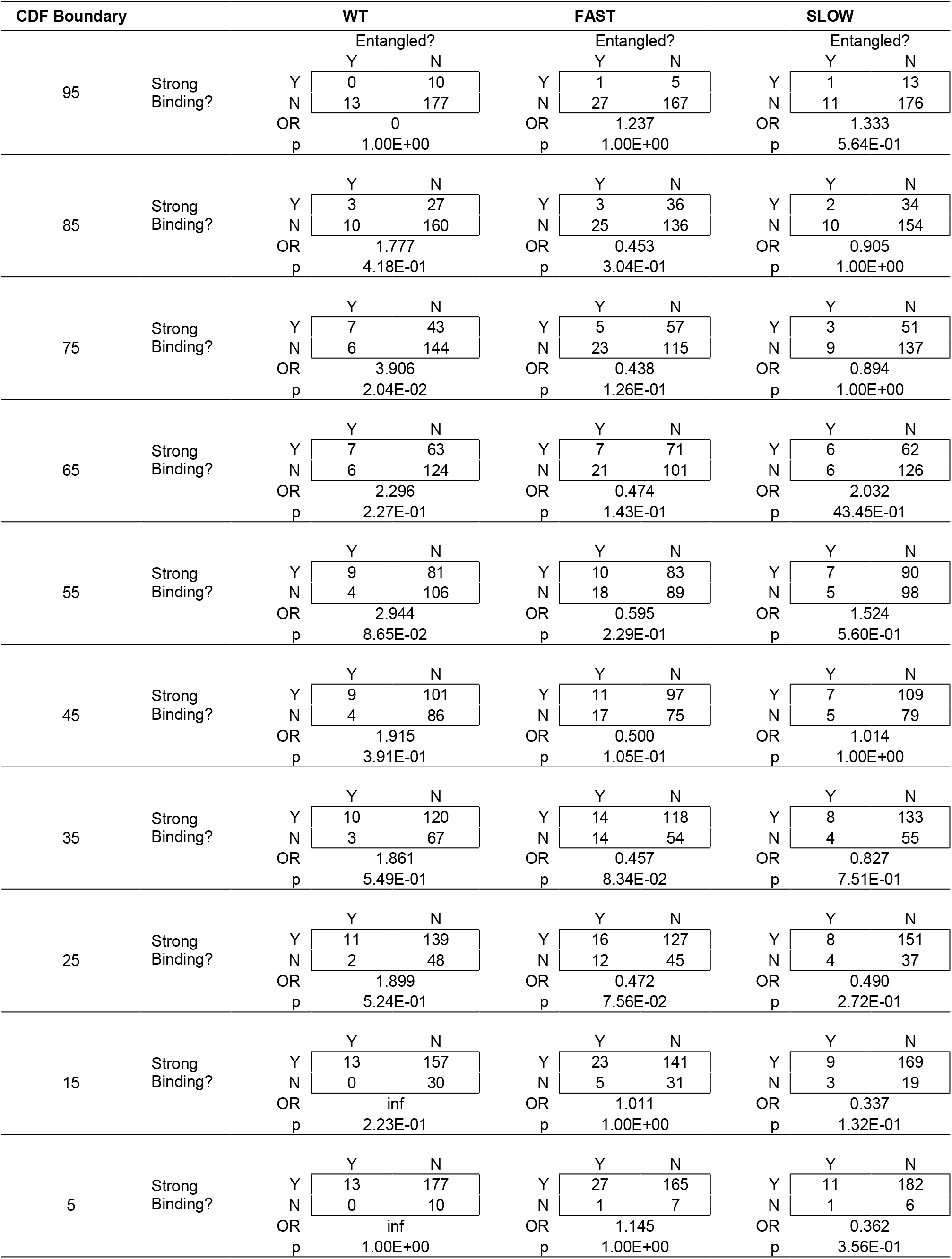
Ribonuclease T entanglement & strong binding region contingency tables

## Notes

### Competing Interest Statement

The authors have declared no competing interest.

